# Evaluating diversionary feeding as a method to resolve conservation conflicts in a recovering ecosystem

**DOI:** 10.1101/2023.11.09.566200

**Authors:** Jack A. Bamber, Kenny Kortland, Chris Sutherland, Ana Payo-Payo, Xavier Lambin

## Abstract

1. The recovery of mammalian predators of conservation concern in Europe is a success story, but their impact on some prey species of conservation concern may cause conservation dilemmas. This calls for effective intervention strategies that mitigate predator impacts without compromising their recovery.

2. We evaluated diversionary feeding as a management intervention tool to reduce depredation on nests of rapidly declining Western capercaillies in Scotland. We studied the influence of diversionary feeding provision on the fates of artificial nests deployed using a replicated and representative randomised landscape-scale experiment. This comprised 30 paired control (no diversionary feeding) and treatment (diversionary feeding applied) sites, 60 in total, each containing six artificial nests distributed across 600 km^2^. The experiment was replicated over two years, and in the second year, the control-treatment pairs were reversed, yielding 60 treatment and 60 control sites and 720 artificial nests.

3. Diversionary feeding substantially reduced depredation of artificial nests, translating into an 83% increase in predicted nest survival over 28 days of incubation. The increase in survival was mostly accounted for by a reduction in the probability that a pine marten, the main nest predator, consumed or cached eggs. Diversionary food also significantly reduced nest predation by badgers, although the magnitude of this effect varied by year.

4. Diversionary feeding is an easily employable method shown in this study to reduce predator impact (functional) without lethal (numerical) intervention. Managers should proceed with its application for conserving capercaillie in Scotland without delay.

## Introduction

Many conservation interventions lack sufficient evidence of effectiveness before implementation despite multiple pressing issues requiring effective intervention due to limited resources and vast demands (Sutherland et al., 2004). This often leads to inefficient and ineffective management that fails to achieve, or worse, is detrimental to, broader conservation goals.

The recovery of mammalian predators across Europe in response to increased legal protection and reduced lethal predator control is a conservation success story. However, recovering predators exploit a range of prey, some of which are of conservation concern or financial interest, leading to conservation conflict (Redpath et al., 2013). Lethal control of generalist predators is widely used as a management intervention to maximise a harvestable surplus or improve declining species’ conservation status (Gibson, 2006). Lethal control is widely accepted when eradicating or controlling damaging non-native predators (Zavaleta et al., 2001). In contrast, lethal control of native predators to protect native species is ethically debatable and has often been shown to be ineffective and poorly implemented, except at small spatial and short temporal scales (Kämmerle & Storch, 2019).

Many generalist predators readily compensate for losses through increased immigration and reproduction, leading to only short-term (i.e., months) reductions in predator density unless control is applied continually (Lieury et al., 2015). Accordingly, meeting the requirements for substantial impact is difficult (Kämmerle & Storch, 2019), as is the lethal control of protected predators, which requires licensing (Sainsbury et al., 2019). Lethal control may, therefore, only extend to a small portion of the predator guild, and the most impactful species may not be affected. Culling also disrupts the predator guild, with consequent changes in the behaviour and density of non-target species (Rees et al., 2023) while disrupting regulatory ecosystem services that some predators provide (Sheehy et al., 2018; Williams et al., 2018). Even for species that are not presently protected, lethal control of predators is not universally deemed ethical by the public (Santiago-Ávila et al., 2018). Therefore, interventions to alter the impact of predation, as opposed to interventions that seek to reduce the abundance of predators, may be more effective for conservation.

The most abundant prey often dominates the diet of mammalian generalist meso-predators, typically small mammals (e.g., voles), but in the face of episodic scarcity of (cyclic) primary prey species, prey switching results in increased predation on alternative prey, such as ground-nesting birds and their nests (Kjellander & Nordström, 2003). In fact, nest loss to generalist predators has been implicated in the decline of multiple ground-nesting birds, including forest grouse and waders (Ewing et al., 2022; Ibáñez-álamo et al., 2015), and there is compelling evidence that ground-nesting seabirds, waders, and gamebird populations can be suppressed by predation (Roos et al. 2018). Specifically, populations of long-lived species with high adult survival and late onset of breeding are more likely to be impacted by predation.

Many ground-nesting birds have adaptations to reduce the impact of nest predation and allow coexistence, such as laying large clutches, re-laying, and camouflage, which reduces variation in population growth rate (Lima, 1987, 2009; Troscianko et al., 2016). However, the level of nest loss a population can withstand depends on its demographic and ecological context (Banks, 1999). For example, if elevated density of generalist predators (e.g. via food subsidy (Pringle et al., 2019) or lack of competitors (Petty et al., 2003)) or more vulnerable nests (e.g. a reduction in safe nesting habitat (Kaasiku et al., 2022) and compounded by deleterious climate and habitat influences (Ibáñez-Álamo et al., 2015) increase nest depredation rates then losses may become additive and a threshold exists below which decline will eventually lead to deterministic extinction and management intervention may be warranted or, in some cases, is the only option.

One promising non-lethal management option is diversionary feeding: the deliberate provisioning of food to change the behaviour of target species and reduce unwanted behaviour (Kubasiewicz et al., 2016) by exploiting the propensity of foraging individuals to exploit the most easily accessed resources (Pyke, 1984). It has been used to reduce the predation impact of single predator species, such as red kites (*Milvus milvus*) on lapwings (*Vanellus vanellus)* (Mason et al., 2021) or kestrels (*Falco tinnunculus*) on little terns (*Sternula albifrons*) (Smart & Amar, 2018).

A notable 22-year-long diversionary feeding trial in a boreal forest landscape (Norway) resulted in population increases in black grouse (*Tetrao tetrix)* and Western capercaillie *(Tetrao urogallus)*, attributed to a reduction in predation by foxes (Finne et al., 2019). In a similar experiment that provisioned foxes with dog food, nest predation during cyclical vole crashes decreased (Lindström et al., 1987). These studies are valuable starting points for understanding how diversionary feeding may influence nest predators specifically. However, the success of diversionary feeding is often species and context-specific (Kubasiewicz et al., 2016). Large-scale experimental evidence is limited yet vital in establishing how diversionary feeding can function as a widely applied conservation intervention and is needed to assess new species-specific contexts and locations or for differing outputs, such as alleviating nest predation pressure.

Forest grouse species (Tetraonidae) are the focus of much interest from a game and conservation management perspective. Culling nest predators is often promoted as a key intervention for grouse population maintenance (Fletcher et al., 2010). One species with a significant conservation focus across Europe is the Western capercaillie. In Scotland, several well-funded conservation initiatives have failed to halt the pronounced decline of capercaillie since the 1970s, with evidence of a further 50% reduction from 2016 to 2020 and extinction being predicted within the next 50 years (Baines & Aebischer, 2023). Climate change is the likely ultimate driver of decline through reduced food sources for chicks and hens (Wegge et al., 2022). However, multiple proximate factors are also implicated in the decline (i.e. fence collisions (Baines & Summers, 1997)), including a significant impact of predation on productivity (Summers et al., 2004).

Lethal control of foxes and crows is common practice across many shooting estates in Scotland, yet capercaillies have disappeared from all but a few shooting estates. The core remnant populations are now found in Speyside in many woodlands where predator control is not carried out. In contrast, two other potential grouse predators, badger (*Meles meles)* and pine marten (*Martes martes,* marten hereafter), are protected species in Scotland and cannot be routinely controlled in the same manner, (MacPherson & Wright, 2021) with the latter being implicated in capercaillie declines (Baines et al., 2016). Lethal control of martens has now been suggested as a possible capercaillie conservation option, resulting in tensions because this would risk undoing conservation gains (marten recovery) whilst also requiring significant scale, effort, and cost to overcome compensation through immigration. Given these legislative restrictions, practical difficulties, a lack of scientific consensus on efficacy, and the intraguild complexities resulting from the disruption of predator communities, there is an urgent need to evaluate alternatives such as diversionary feeding. Particularly given its potential to influence the behaviour of multiple predators.

Considering that evidence on the effectiveness and practicalities of diversionary feeding has been mixed, we respond to a need for experimental evaluation of diversionary feeding, evaluating its potential application as a practical and feasible management intervention to decrease nest predation. We do this through large-scale, experimental deployment of diversionary feeding coupled with control sites with no feeding. Specifically, the focus was on the protected marten as an important nest predator and the critically endangered capercaillie. Our experiment compared artificial nest survival in a control-treatment design to evaluate how diversionary feeding influences the rate at which nests are depredated. Our experimental approach provides a robust, accurate, and comparable index of predation change, with nest failure purely being related to predation pressure and not alternative factors, such as nest abandonment due to adverse weather, as may be true with real nests.

## Materials and Methods

### 2.1 Study Area

This study was conducted in the Cairngorms Connect landscape (FLS, Wildland Ltd, RSPB, Naturescot), a 600km^2^ ecological restoration project on the western side of the Cairngorms National Park, Scotland (57°09’47.5“N 3°42’47.0” W, Figure 1). The landscape consists of remnant Caledonian and plantation pine forests (mainly *Pinus sylvestris*), with a mixture of bogs, heaths, and some deciduous woodlands. Management includes intense culling of *Cervidae* (red and roe deer) to allow forest regeneration, and, unlike in more traditional neighbouring estates, there is no control of predators. The area encompasses the core of the remaining population of Scottish capercaillie (Baines & Aebischer, 2023). The predator community includes badger, fox, marten, carrion crow (*Corvus corone)*, common buzzard (*Buteo buteo)*, and ten scarcer raptor species. Field and bank vole numbers, assessed bi-annually via the vole sign index respectively (Lambin et al., 2000) and live trapping with the small quadrat design (Myllymaki et al. 1971), were low, including a crash (2021) and early increase (2022) years of a 3–4-year population cycle. Accordingly, baseline nest predation rates were expected to be high.

**Figure 1.**
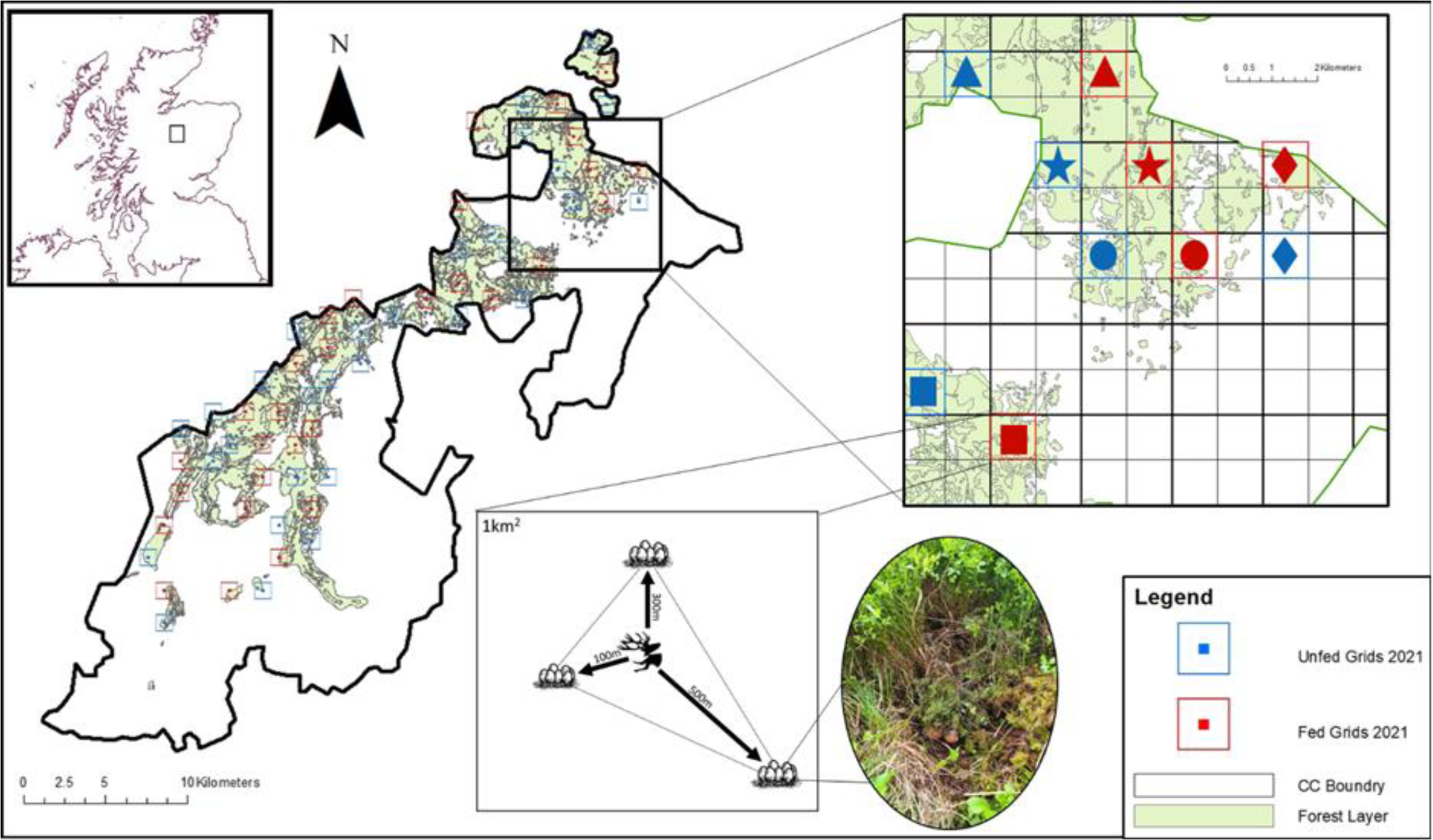
Illustration of Experimental Design. The Main map shows the forested areas of the Cairngorms connect landscape, which was our scope of inference. The red and blue squares show the locations of our 60 sampling grids, with feeding sites (2021) in red and control sites(2021) in blue. The highlighted rejoin in the top right zooms into our sampling grid to show the site structure of 1 km2 sampling grids; “paired” sites are shown with matching shapes. The lower 1km^2^ area shows an example of the internal structure of a diversionary feeding site, an example of an initial deployment of nests and a central feeding station. Control sites mimicked this structure without the feeding station. An exemplary artificial nest with heather cover is also included.

The study area was selected in response to land managers in the Cairngorms Connect land partnership having ceased lethal predator control and, having been inspired by the findings of Finne et al. (2019), being eager to rigorously assess this non-lethal method whilst being deployed as capercaillie emergency conservation intervention. The Cairngorms Connect board deemed the risk of adverse outcomes acceptable given the wide availability of deer entrails in the area, and the experiment largely redistributed this food in time and space.

### 2.2 Experimental Design

We performed a randomised landscape-scale experiment with paired control (unfed) and treatment (fed) sites swapped between years. Defining the site as 1km^2^ square grid cells, we deployed 60 pairs (30 control and 30 treatment) each year across two years restricted to forested areas (National Forest Index, min of 1.7 hectares of forest cover). The size of the grids was chosen to encompass the typical daily home range of a marten (49 ha in females and 54 ha in males (Zalewski et al., 2004)). Diversionary feeding treatment was randomly assigned to all but 4 of the 30 cells, and a second control cell was selected to be at least 1km^2^ (edge-to-edge) from any diversionary feeding cell to maximise treatment independence while maintaining the pairing (Figure 1). The centre of treatment cells had a feeding station (see below), and each grid cell contained three artificial nests at once across two deployments per year, totalling six (see artificial nests below). Due to constraints resulting from the convoluted shape of the study area, 8 cells with edges closer than 1 km were given the same treatment to maintain the independence of treatment. The experiment was conducted over nine weeks between 24^th^ April and 1^st^ July 2021 and 2022, coinciding with tetraonids’ nesting, re-nesting, and early brooding periods (Summers et al., 2004). Diversionary feeding and control treatments were swapped within pairs between years, further reducing any influence of cell properties beyond randomisation.

### 2.3 Diversionary Feeding Stations

To maximise the applied relevance of the experiment, we provided supplementary by-products from ongoing deer culling. This is a resource known to be consumed by predators when left in situ as gralloch (organs left after culling) and carcasses, which was more cost-effective than provisioning other food sources. The nine-week timeframe of provisioning was deliberately short to avoid any numerical predator response, a potential risk with diversionary feeding (Kubasiewicz et al., 2016). The feeding period also coincides with the time of more abundant food (Spring), meaning that diversionary food is more likely to provide an alternative, not a supplement.

Deployment of diversionary feeding was timed close to egg laying to avoid any increase in predator abundance within the vicinity of feeding stations after territories and breeding decisions of predators were likely fixed with marten delayed implantation occurring in Feb-Apr (Jonkel and Weckwerth, 1963). Feeding stations were deployed within ∼100m of the centre point of grid cells and were replenished every two weeks with ∼10kg of deer carrion (for 8 weeks); the weights of remaining food were recorded with a spring scale to monitor depletion. Non-consumed food was left in situ to increase scent cues. Replenishment ensured food was always available, even if predators found decaying meat unappealing (Moleón and Sánchez-Zapata, 2021).”

All feeding stations were monitored using remote camera traps (Browning Recon Force Advantage model: BTC-7A), set to record three-shot bursts with a five-second interval between captures to establish uptake of diversionary feeding treatment by target and non-target species. Any movement of food upon visits to sites was recorded.

### 2.4 Artificial Nests

We used the fate of artificial nests rather than nests of wild capercaillie and black grouse as the response variable owing to the scarcity of these focal prey and to minimise disturbance. Nest deployment within the sample sites was limited to daylight hours to limit any disturbance to capercaillie in the area; no capercaillies were flushed from their nest. Approximately 7 days after establishing the feeding sites, we constructed 3 artificial nests, containing 7 eggs, in each control and treatment grid (N=180 nests). The three nests were placed 100m, 300m and 500m from the centre of the cell. Nests were made to resemble capercaillie nests: a shallow depression at the base of a tree filled with plant material, covered with dwarf bushes twigs to mimic the visual camouflage an incubating hen provides. We ensured that at least 1.5 eggs were visible to an observer standing 3-5 m away from the nest to allow for nest detection by a visual (avian) predator. Each artificial nest had six small domestic hen (*Gallus gallus domesticus)* eggs, resembling capercaillie eggs in size and colour (Mortola & Al Awam, 2010; Rosenberger et al., 2017). A 7^th^ egg was drained and filled with “Parasoy” soy wax blend, tethered to the ground, to aid in identifying predators through tooth and bill marks. To reduce human scent that may affect discovery rates (Weldon, 2021), eggs were stored on pheasant feathers for seven days before deployment, rubber gloves were worn when handling eggs, and rubber boots and field clothes worn during deployment were stored outdoors.

A second deployment of nests, mimicking re-laying, occurred at either date when predation was detected or the hatch date of the first deployment, at the same distance interval from the cell centre but 50-100 m away from the previous nest to avoid predator bias. Secondary nests contained only 3 hen eggs plus one wax egg, mimicking the lower clutch size seen in relaying (Storaas et al., 2000) and were not replaced once depredated or if they survived 28 days. Three hundred and sixty nests were deployed each year (720 total).

Nests were checked every 14 days, with two visits spanning the 28-day capercaillie incubation. Checks were conducted visually from 3-5 m away. The location of the artificial nests at each distance was determined using a randomly generated compass bearing from the centre of the grid at the specified distance.

If a nest site was disturbed or the 1.5 visible eggs could not be seen during nest checks, the nest was inspected in more detail. Depredation was deemed to have occurred if any hen egg was damaged or removed. Field signs, including marks on wax eggs and patterns of nest disturbance, were recorded to ascertain which predator was likely responsible (Summers et al., 2004). The assumed predator was assigned for each depredation event before checking camera trap data to allow unbiased validation.

In addition to the 14-day visual checks, camera traps (see model above) were positioned at a subset of artificial nests set to record 10-second-long videos and distributed equally, but randomly, between treatments and distances. Thus, 10 nests per distance from the cell centre had a camera for each treatment (60 per treatment, totalling 120 per year). This allowed the specific identification of nest predators and was used to validate field interpretation of wax egg markings and nest signs.

### Data extraction

Camera trap photos taken at feeding stations were identified at the species level using the metadata tagging software DigiKam 7.3.0, following the ‘CamtrapR’ workflow (Niedballa et al., 2016). Videos taken at artificial nests were viewed, and species-specific detection histories were generated manually by recording the time and date of a depredation event. The assumed responsible predator of depredated nests for nests without cameras was inferred from field signs. Video footage from camera traps allowed validation of assumed predators from signs with confirmed predators on camera traps, showing a 97% (106 of 109) success rate in correctly identifying nest predators when an assumed predator was assigned (See Appendix 1). We collected the fate of 720 nests; 15 nests were excluded from specific fate analysis, known to have been depredated by a fox (n=2), corvids (n=7), or rodent (n=6) owing to low sample size and 28 nests depredated by non-identified species as they likely formed a heterogeneous group (12 Treatment: 31 Control).

### Statistical analysis

We modelled the fates of artificial nests using multinomial logistic regression with three possible fates:

1) 28 days, 2) depredated by marten, and 3) depredated by badger. We analysed the multinomial responses using covariates that reflected the experimental design: diversionary feeding treatment to quantify the effect of diversionary feeding, distance from the grid centre/feeding station to quantify any spatial decay of the feeding effect, and interaction between distance and treatment. We also included a year as a fixed effect for annual variation, e.g., in field vole prey abundance. We added grid cell identity (n=60) as a random effect to account for the non-independence of nests within a grid resulting from similarities in local influences on nest predation, such as local predator abundance and habitat, across the two years of study.

Multinomial models were implemented using a generalised additive random effects model (GAM) in package ‘mgcv’ (Wood, 2017). We used 1000 simulations from the model using the function “Predict” to produce estimates of the marginal (population-level) probability of each fate and associated 97.5% confidence intervals (CI). All statistical analysis was performed using R (version 4.1.3).

## Results

### Uptake and usage of feeding stations

On average, 57.5kg (range: 38 - 81 kg) of deer meat was deployed per feeding station each year, with an average of 10.5kg (range: 6 - 14 kg) added every two weeks according to the weighed assessment of depletion at restocking visits.

Over 340,000 photographs were collected from the 60 feeding stations across 3,912 camera trap days. They include 142,179 images of potential egg predators recorded in 3,726 independent visits. Martens accounted for 13.0% of these visits (19,374 images, 486 detections), badgers for 22.5% (54,024 images, 839 detections), and foxes for 10.3% (12,946 images, 383 detections). We also detected avian nest predators: 35% of the visits were by raptor sp. (buzzard, golden eagle, red kite (31,837 images; 1306, detections)) and 17.5% by corvids (jay, crow (20,410 images; 639 detections). Fifty-four of the 60 feeding stations were visited by predators (86.6% in 2021, 93.3% in 2022). Badgers, foxes, and buzzards visited similar proportions of feeding stations (58%, 57%, and 55%, respectively), followed by martens (43.33 %) (See Appendix 2). As predators had moved deer carrion away from the camera fields of view at 132 restocking visits on 240 instances, it was not possible to estimate the amount of diversionary food consumed by each species.

### Nest fate with diversionary feeding

Forty-nine percent of artificial nests (353/720) survived the full 28 days, meaning that 51% of nests experienced depredation. Recorded nest predators were martens (38%, 268 cases), badgers (6.9%, 50 cases), and other species (unknown, fox, corvid, and rodent (6.8%, 49 cases)). However, fewer nests survived in control sites (128, 35.5%) than in diversionary feeding sites (228, 63.3%), reflecting fewer nests depredated by martens (Control: 170, Treatment: 98) and badgers (Control: 31, Treatment 19).

Multinomial logistic regression revealed that nest fates were associated with two experimental variables: treatment and year, but in different ways (Table 1). There was no effect of distance on the expected fate of a nest in either control or treatment sites or between species. The probability of marten depredation was substantially lower in a treatment site compared to a control site (−1.494, SE 0.309, p<0.000), and this did not vary between years (−0.007, SE 0.172, p= 0.969). This amounts to marten predation probabilities of 0.22 (CI 0.151-0.318) and 0.52 (CI 0.40 0.64) in fed and unfed sites, respectively (Figure 2). The probability of badger depredation was also significantly reduced by diversionary feeding (−1.723 SE 0.622, p=0.006), with a significant additive influence of year reflecting higher badger depredation in 2022 (0.694 SE 0.325, P=0.033). This provided predicted changes to fate badger of 0.085 (CI 0.0187-0.227) to 0.03 (0.005-0.098) in 2021 and 0.15 (CI 0.04-0.366) to 0.058 (CI 0.011-0.175) for control and treatment respectively (See Figure 2). Considering this change in the probability of predation fates, the predicted probability of nests surviving changed from 0.406 (CI 0.303-0.523) in control up to 0.744 (CI 0.645-0.828) with treatment, an increase of 82.5% (See Figure 2). This change occurs mainly due to the change in the predicted probability of marten depredation, reducing in value with diversionary food provision.

**Figure 2.**
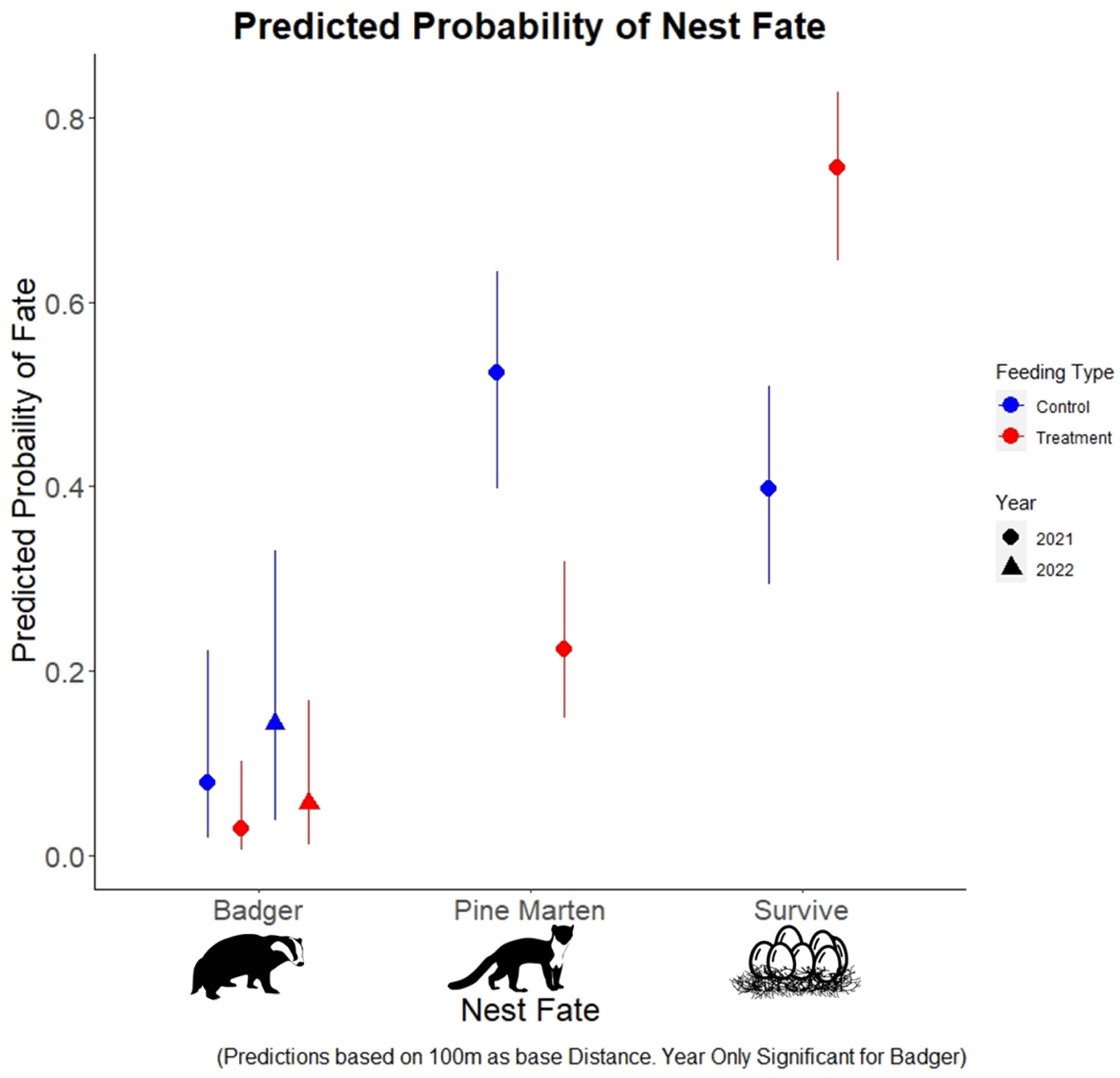
Multinomial logistic regression showing predicted probabilities of nest fate. Predictions are made for a baseline of 100m for all three fates badger, pine marten and survived. We present predictions for badger for both 2021(●) and 2022 (▴), as it was highlighted to be significantly different by multinomial modelling. Error bars show 97.5% confidence intervals for control (black) and treatment (red).

**Table 1.**
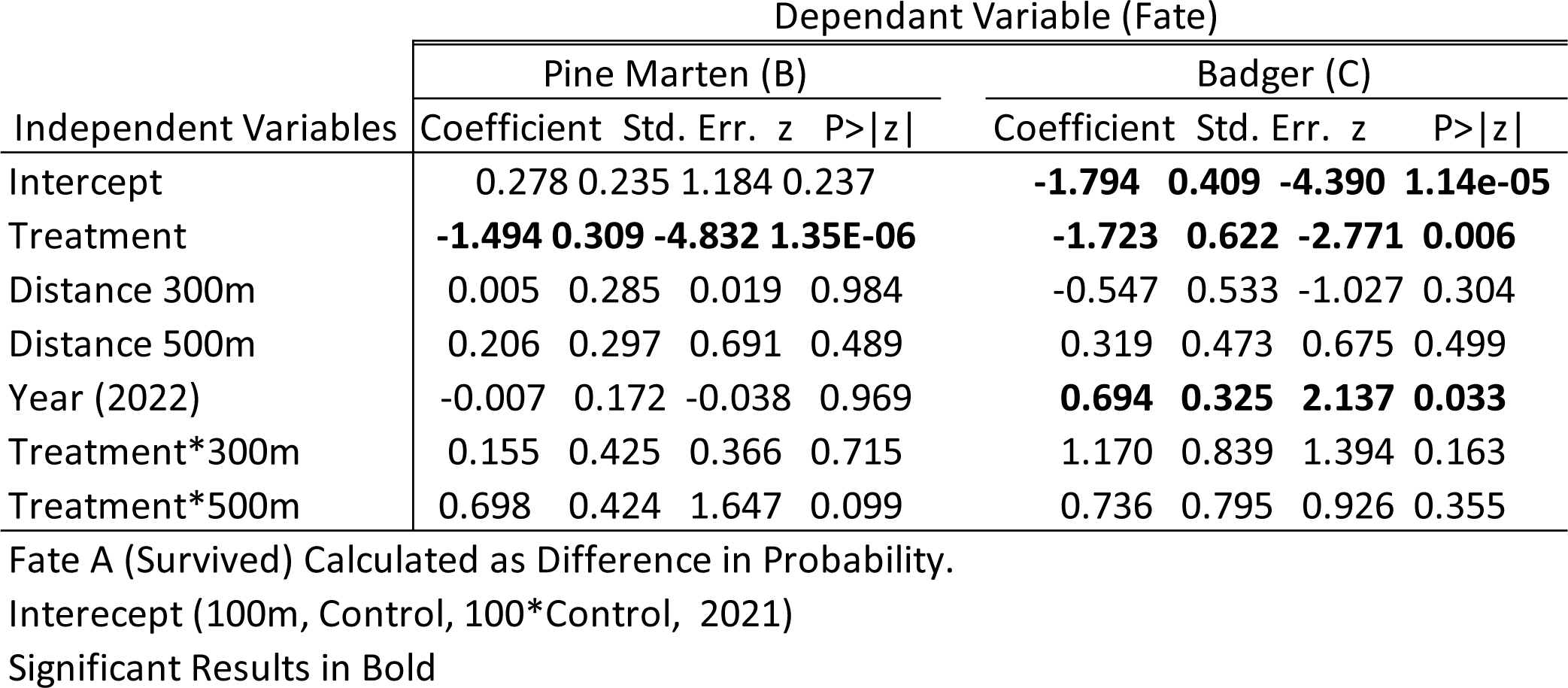
Coefficients of multinomial logistic regression, showing changes in the log probability of the fate of artificial nests. The reference fate (or base fate) for the model is that a nest survived for 28 days, with the changes shown with fate B and C being the change in odds of being fate “survived”. Significant results are highlighted in bold.

These predictions align with survival analysis for all nests, without consideration of fate (See Appendix 3).

## Discussion

We evaluated how diversionary feeding alters the rates of nest depredation by meso-carnivores in a boreal forest landscape. We found that diversionary feeding almost halved depredation rates of pine martens and badgers on artificial nests over the 28-day incubation period. Predicted nest survival probability increased by 82.5% (from 0.406 to 0.733) with the provision of diversionary feeding. Using a fully randomised, well-replicated, landscape-scale experiment allows us to infer causality between short-term provisioning of diversionary feeding and reduced depredation of artificial nest predation. Should they extrapolate to real capercaillie nests, our results present diversionary feeding as a viable non-lethal option for reducing nest predation on ground-nesting birds of conservation concern across the boreal zone, providing a form of control of the impact of predator presence without sacrificing ecosystem benefits, or losing public support. The presence of a validated alternative to lethal intervention raises questions as to whether practitioners have the social licence to cull one protected species for the protection of another.

### Intervention for Protected Predators

Diversionary feeding almost halved both marten and badger depredation rates but from different baselines. Because of these two species’ reduction in nest depredation, artificial nest survival increased by 83% (Figure 2). The risk of depredation by pine marten was five times higher than by badgers. Hence, the reduction of pine marten impact is proportionally greater for nest survival. This is despite martens having confirmed access at only 43% of feeding stations from cameras. This could be due to the localised redistribution of diversionary food into the area surrounding feeding stations by other predators, with large pieces of carrion found up to 50m away from feeding stations, causing imperfect detection. Similar food redistribution around carcasses by badgers and foxes has been seen at a maximum of 103m (Young et al., 2015). Whilst the separation distance of treatment and control sites encompassed the daily movement patterns of pine marten, full home ranges may have been larger for some individuals. This may mean that they may have used multiple sites, however, this would likely have lowered any effect of treatment.

Both badger and marten numbers have been suppressed historically due to persecution. Pine marten were driven to localised extinction but have recovered since the first re-sightings in the study area in 1994. It has been suggested that this recovery and the perceived high density of pine martens may be the reason for unsustainably high nest predation rates (Baines, Aebischer, & Macleod, 2016). Surveys in 2012 within the research area estimated densities of 0.07 - 0.38 individuals per km^2^ using spatially explicit capture-recapture of non-invasively collected hair in sites within our research area (Kubasiewicz et al., 2017). A 2020 survey using similar field and analytical methods indicates no rise in density Hobson et al. (2023). These values are around the lower density estimates for pine marten elsewhere, such as Białowieża Forest, ranging from 0.363-0.757 individuals per km^2^ (Zalewski and Jedrzejewski, 2006). This indicates that modifying marten feeding behaviour through diversionary feeding is a realistic, sustainable, and evidence-based alternative to lethal control, especially if marten populations are not above normal densities. Future studies on the efficacy of this method where marten densities are higher would add to the evidence supporting the method for intervention where marten numbers are abnormal.

Land manager perceptions (before this experiment) were that badger numbers have increased in the study area; while no formal estimates have been presented, we found that badgers were widely distributed, accessing 58% of feeding stations and depredating artificial nests in a pine forest, where historically, they were deemed mostly absent. This could reflect a recovery by badgers and another reinstated source of predation pressure that could also be halved by diversionary feeding.

### Interpreting artificial nest data

There are known caveats to interpreting the absolute depredation rates on artificial nests and necessary conditions for interpreting differences in predation rates in an experimental context such as ours. To avoid overemphasising some predators’ impact and underestimating others’ impact, the predators responsible for actual nest predation must match those of actual nests (Pärt & Wretenberg, 2002). For marten, the main predator in this study, we can be confident that changes seen in nest predation may translate to actual nests, as they have been seen to predate actual nests at a similarly high rate in other studies. Summers et al. (2009) sampled actual capercaillie nests in the same region with camera traps (N=22); in this instance, all confirmed losses were due to marten predation.

Consideration of other studies on predation rates on capercaillie nests reveals that the fate of artificial nests in this study aligns with what is seen elsewhere in their range. Predation rates of capercaillie nests in a stable population in southern Norway ranged from 48 to 90% according to the stage of the vole cycles (Wegge & Storaas, 1990). In Scotland, observed predation rates revealed by camera traps deployed on actual nests ranged between 42 and 68% (Summers et al., 2009). Our estimates of 65% predation rates on our artificial nests in control sites are within these ranges, with the observed 37% in the presence of diversionary feeding as low as the lowest value seen in Norway in a peak vole year. Thus, present nest predation rates in Scotland are not abnormally high, especially considering our study took place during low vole years when predation rates were expected to be at the high end of observed rates.

In this study, fox and corvid depredation of artificial nests was lower than on actual nests studied by Baines et al. (2004) in the same region. The presence of camera traps at some, but not all, false nests may have deterred foxes, given the prevailing persecution (Zalewska et al., 2021). Conversely, our efforts to reduce human activity at our artificial nests may have precluded the inflated corvid predation that often occurs where humans interact with nests (such as trails and markers (Picozzi, 1975)). While some raptor species influence red grouse recruitment, the impact occurs through chicks, not nest predation (Thirgood et al., 2000), so our results are not unexpected.

### Consideration of risk factors

The duration of our experiment, from late April to early July, was chosen to reduce the risk of a numerical response through aggregation by predators potentially subsidised by deer carrion. Predator territories and the number of embryos were long established before the deployment of diversionary feeding. Thus, we infer diversionary feeding changed predator foraging (functional response), not numbers (numerical response). However, deer culling activities mainly occur in winter in Scotland. It is not unlikely that predators’ numbers are elevated by the overwinter provision of gralloch (when it is not removed; 23% of the research area) across the landscape, as was the case before our experiments.

Increased nest survival may not directly translate to more chicks reaching adulthood, hence productivity (Saniga, 2002). If decreased nest predation and increased chicks make a breeding area more attractive to predators aggregating in areas with diversionary feeding, this may elevate the predation rate (Pakanen et al., 2022). However, as capercaillie chick biomass is tiny relative to all other prey exploited by pine martens, badgers and other mesopredators, we view this form of compensatory increase in predation to be unlikely but worthy of further study.

### Management Implications

Using one fieldworker, this experiment covered most of Scotland’s core residual range of capercaillie. Deployment of 30 feeding stations across five land ownership areas took five days. The focus on good experimental design for robust inference meant that deployment was labour-intensive; practical deployment would likely be easier, with no need for strict separation and designation of “control” and “treatment” sites. This would also allow for sites to be deployed at a higher density within the areas needed for intervention. Our design with nests at three distances from the food dump found no clear evidence that the depredation rate changed significantly with increasing distance from the feeding station up to 500m. Based on the study design, the minimum effective range of influence of one station per km^2^ of suitable habitat was shown in this study. However, with no significant reduction seen in efficacy within that distance, the spacing could be more comprehensive for practical deployment. Logically, the influence of feeding on nest predation rate must reduce eventually, as seen with the minimum 1.5 km^2^ separation of feeding stations to control site nests. Indicating that 1 feeding station for 1.5 km^2^ of capercaillie habitat would be suitable. Using by-products from existing deer culling efforts meant the cost of providing food was low and may even have reduced the disposal cost. Based on the total food deployed at each feeding station across our experiment, at maximum, a feeding station would require approximately 80kg of carrion, which is the equivalent of an adult male red deer (Reby & McComb, 2003). With large numbers of deer being culled, there is no limitation to the amount of deer viscera that could be made available by real-time supply or freezing byproducts during peak cull periods. Thus, there is little doubt that diversionary feeding could be rolled out across the remaining range at little cost with potentially substantial benefits to a ground-nesting species in decline. To establish if a reduction in nest predation alone can lead to capercaillie recovery in the face of climate change, monitoring the influence of diversionary feeding should be performed, emphasising evaluating the full impact on productivity. While our protocol strove to avoid increasing the abundance of predators, it remains essential to establish if this was effective. Evaluating whether supplementary food or diversionary might inflate mesopredator density over time, e.g. through increased offspring survival, alongside diversionary feeding, is an area for future research.

The observed substantial reduction in artificial nest depredation demonstrates the potential of diversionary feeding as an effective non-lethal intervention for conserving ground-nesting birds. It can be used with protected and recovering predators of conservation concern, hence alleviating conservation conflicts. Particularly, given pine marten recolonisation following legal protection, implementing non-lethal management action that mitigates the impact of predation is feasible and supported by the evidence presented here. No major obstacles should exist to implementing diversionary feeding whilst further monitoring impacts throughout the historical range of capercaillie now shared with the native pine marten.

## Acknowledgements

We acknowledge funding from NERC SUPER DTP, reference NE/S007342/1 and Forestry and Land Scotland to JB. We thank all Cairngorms Connect partner organisations, funders and staff for in-kind contributions, including: Bobby Innes, Cieran Watson, Tom Cameron from Forestry and Land Scotland; Anders Poulsen, Thomas MacDonell, Davie McGibbon, Donnie Ross, Ronan Dugan, Grant Ashley, Marc Willis from Wildland Ltd; Richard Mason, Michael Butler, Cameron Waite, Fraser Cormack, Jack Ward from the RSPB; Endangered Landscape Project, and Elsa Fleurial and Titouan Sanglard for assistance.

## Appendix Document

## Appendix 1 Nest Camera Trapping Assumed and Confirmed

Comparison of assumed predators from signs with confirmed predators on camera traps showed a high success rate in correctly identifying nest predators when an assumption was made. In many cases where an observer was incorrect in assigning an assumed predator (pine marten only), the pine marten was the primary nest predator. Still, the nest site was visited afterwards by a secondary predator. When assigning an unknown mammal, in some instances, it was due to multiple predators interacting with the nest site before the observer could look at the predation signs, changing the indications, or, in the case of the fox, having so few instances of occurrence, signs were not clear.

Out of 115 nest sites predated by predators with a camera trap deployed on them, 106 were able to detect the true nest predator successfully, and six failed to detect any interaction with the nest due to camera trap failure. Overall, observers were correct over 99% of the time when assigning an assumed predator, or 97% when considering the assignment of unknown to two nests later confirmed to be a fox. This gave us confidence to use assumed and confirmed predators in further fate analysis.

**Table 1.** Fates of nests of camera-trapped sites, with confirmed predation. The total number of assumed assignments is compared with the number of confirmed predators and if the assignment was correct. In the cases of incorrect assignment, the true predator was highlighted in “Incorrect classification”. In the case of unknown mammals, the incorrect classification highlights who the truly responsible predator was.

| Comparison of Assumed Accuracy (Nests with Camera Traps) |  |  |  |  |  |
| --- | --- | --- | --- | --- | --- |
| Predator | Assumed Assignments (Total) | Number Confirmed | Correct Assignments | Accuracy (%) | Incorrect Classification |
| Pine Marten | 72 | 72 | 70 | 97.2 | 1 Marten & Fox, 1 Marten & Badger |
| Badger | 18 | 18 | 18 | 100 | NA |
| Fox | 0 | 2 | 0 | NA | NA |
| Rodent | 4 | 4 | 4 | 100 | NA |
| Avian | 0 | 0 |  | NA | NA |
| Unknown_Mammal | 13 | NA | NA | NA | 1 Badger, 8 Pine Marten, 2 Fox, 1 Dog & Pine Marten, 1 Badger & Pine Marten. |
| N=109 (6 Camera traps failed to detect a predator) |  |  |  |  |  |

## Appendix 2 Feeding Station Access

Feeding stations were monitored via camera traps across the entire sampling regime; camera images were tagged using the methodology described in the main text across the 9 weeks of camera trap monitoring of diversionary feeding. We highlight below the minimum number of feeding sites with at least 1 detection of a feeding predator. The minimum is due to a large amount of food dispersal from the central monitored point preventing perfect detection. Raptor species were grouped into one category to include rare detections, such as White-tailed eagles, but mostly were detections of buzzards. Domestic dogs were accidentally detected at 25% of feeding sites, with them not being kept on leads as is advised during the breeding season. Further breakdown of key predators can be found in the main text.

**Figure 2.**
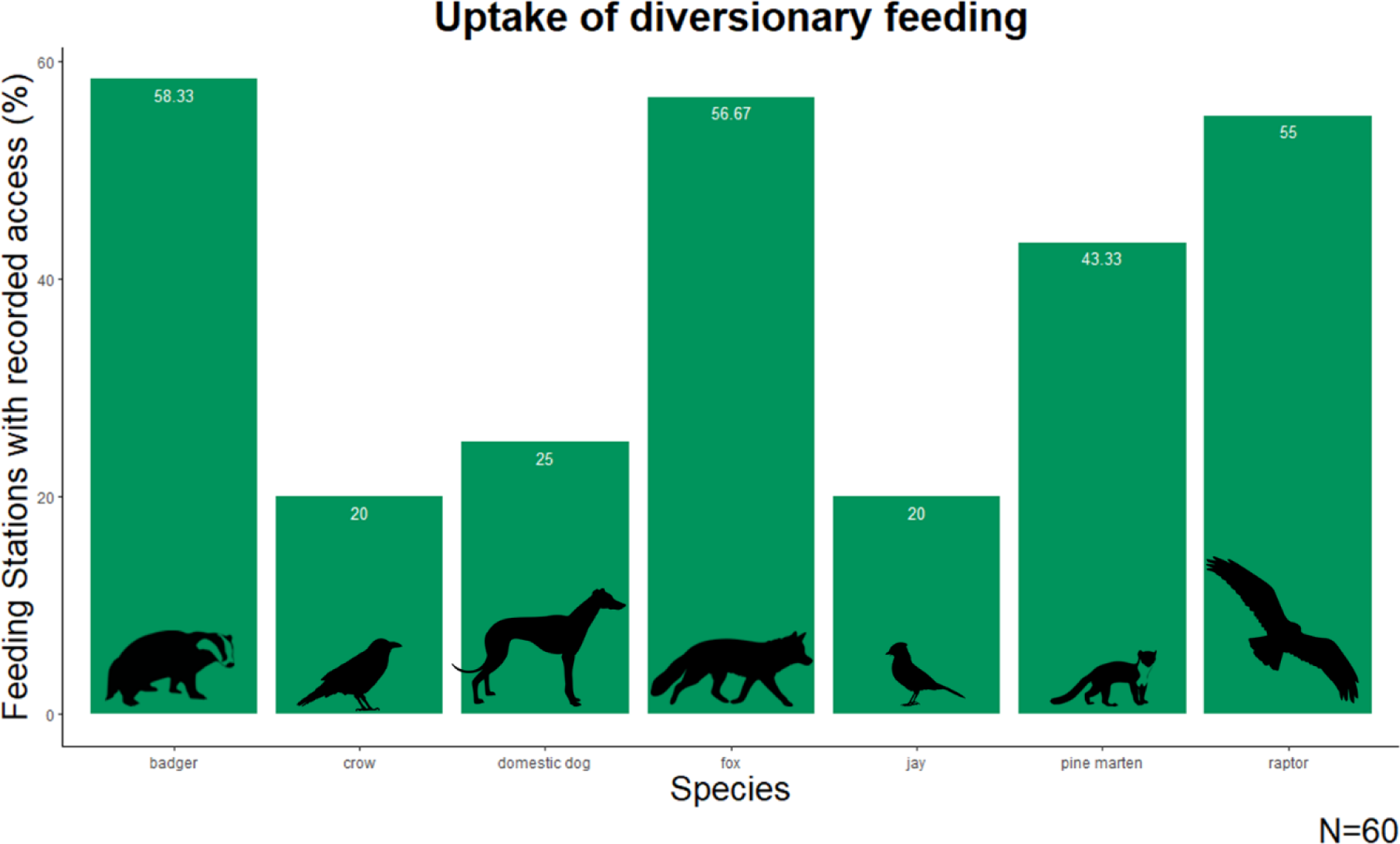
Shows the proportion of feeding stations with confirmed feeding by predatory species via camera trapping. The y-axis shows the percentage of all 60 sites that had a detection of each of the species listed on the X-axis.

## Appendix 3 Survival Irrespective of Fate Kaplan Meir

Primary Exploration of false nests was done via Kaplan Meir; this looked at the entire 720 nests in the study without any information based on the assumed fate.

The daily status of each artificial nest over 28 days was coded as a string of 1 (intact) and 0 (depredated) using the known predation date for nests with cameras and the mid-point between visual checks for depredated nests without cameras. We estimated the influence of treatment by fitting the Kaplan-Meier survival probability curve using the ‘survival’ package (Therneau *et al.,* 2023). This allowed us to assess baseline fate (survived or predated) for all 720 nests over the 28-day sampling period. Large numbers of deaths occur on day 7 and day 21 due to the assumption of a central death date of nests between 14 and 28-day nest checks. These large dips indicate the trapping regime, not actual deaths. Links with broader multinomial analysis are made in the main text.

When assessing the fate of all nests (N=720), irrespective of fate via Kaplan meir, all nests also showed an increase in survival with treatment. Base survival of nests (Survived vs predated) showed an estimated control survival of 0.345 (CI 0.281-0.422); with the provision of diversionary feeding, this increases to 0.650 (CI 0.585-0.723), across 28 days. This is an 88% increase, providing confidence in the accuracy of our multinomial predictions.

**Figure 3.**
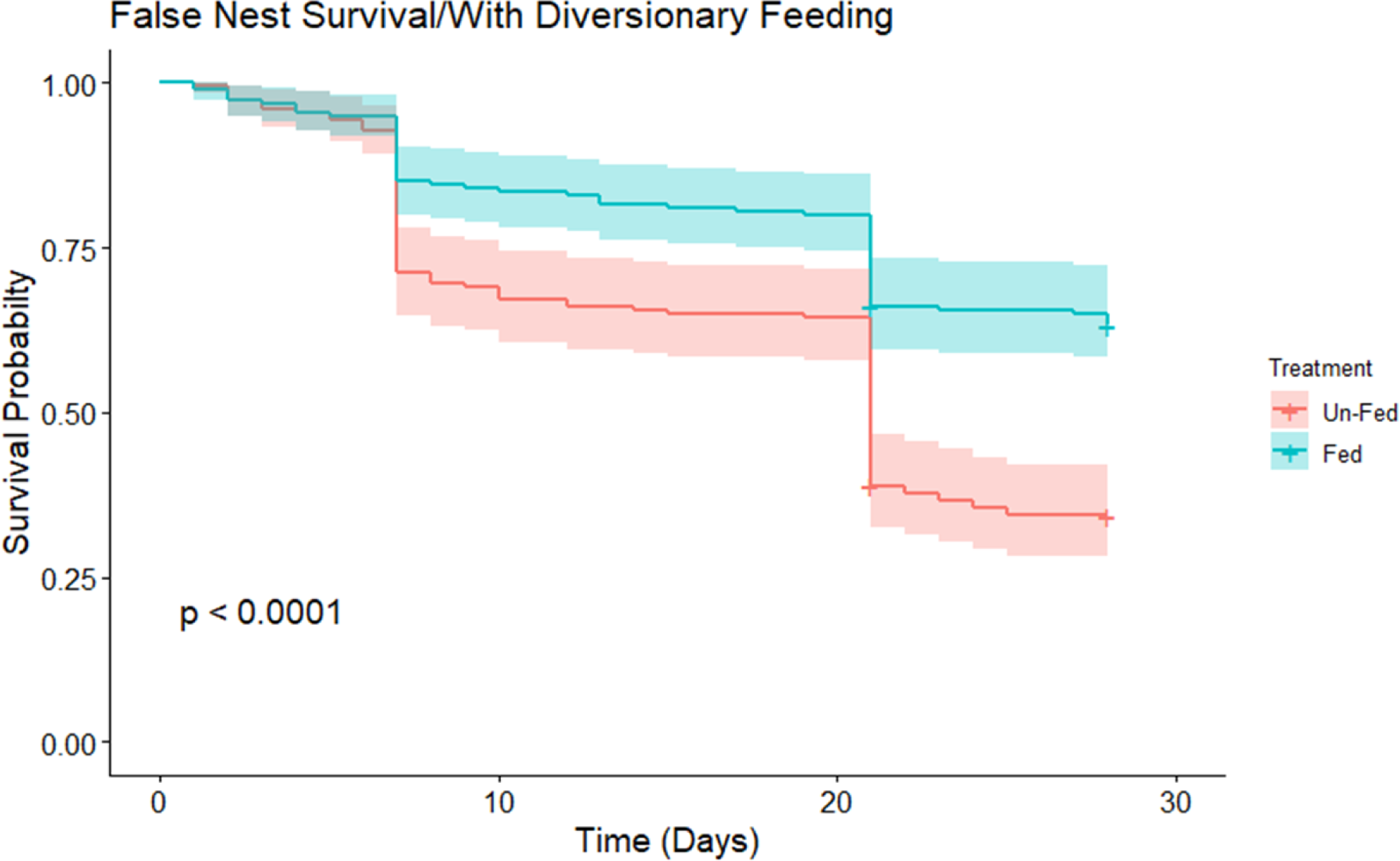
Kaplan Meir survival curve visualising artificial nest survival over 28 days for control and diversionary feeding (legend) diversionary feeding (N=720). Survival across our research period is visualised using Kaplan Meier across 28 days of life for each nest; check days at 14 and 28 days are shown with central death dates at days 7 and 21.

